# An artificial neural network identifies glyphosate-impacted brackish communities based on 16S rRNA amplicon MiSeq read counts

**DOI:** 10.1101/711309

**Authors:** René Janßen, Jakob Zabel, Uwe von Lukas, Matthias Labrenz

## Abstract

Artificial neural networks can be trained on complex data sets to detect, predict, or model specific aspects. Aim of this study was to train an artificial neural network to support environmental monitoring efforts in case of a contamination event by detecting induced changes towards the microbial communities. The neural net was trained on taxonomic cluster count tables obtained via next-generation amplicon sequencing of water column samples originating from a lab microcosm incubation experiment conducted over 140 days to determine the effects of the herbicide glyphosate on succession within brackish-water microbial communities. Glyphosate-treated assemblages were classified correctly; a subsetting approach identified the clusters primarily responsible for this, permitting the reduction of input features. This study demonstrates the potential of artificial neural networks to predict indicator species in cases of glyphosate contamination. The results could empower the development of environmental monitoring strategies with applications limited to neither glyphosate nor amplicon sequence data.

**Highlight bullet points:**

- An artificial neural net was able to identify glyphosate-affected microbial community assemblages based on next generation sequencing data
- Decision-relevant taxonomic clusters can be identified by a stochastically subsetting approach
- Just a fraction of present clusters is needed for classification
- Filtering of input data improves classification

## Introduction

Monitoring the environmental status of the Baltic Sea is required by law as part of the HELCOM agreement (Backer et al., 2010). This includes distinct events such as contamination. In this study, glyphosate as a potentially harmful herbicide (Van Bruggen et al., 2018) as well as a phosphorus-providing substrate (Hove-Jensen et al., 2014) was used as a model contaminant to investigate novel means of monitoring. Glyphosate has been the most-applied herbicide globally since the 1970s. Recent studies have proven that glyphosate is mobile despite its soil adsorption characteristics (Bergström et al., 2011; Kwiatkowska et al., 2016; Myers et al., 2016). Due to its intensive use in agriculture around the world, glyphosate is present in significant quantities in soil and groundwater (Battaglin et al., 2014) just as it enters the Baltic Sea (Skeff et al., 2015). A lab microcosm experiment was set up in which the herbicide was added as a stressor to a brackish-water microbial community. To assess the impact of glyphosate independently of specific glyphosate detection methods, a combined approach of artificial neural networks (ANNs) and next generation sequencing (NGS) technologies was applied. The goal was to determine whether altered microbial community compositions – classified by the ANN - can serve as a marker of a glyphosate disturbance. An ANN was trained on compositions, which were declared as glyphosate-impacted or not. The ANN was then challenged to classify a previously unknown sample with regard to the presence of glyphosate.

Machine learning algorithms and in particular artificial neural networks are important tools in data analysis and decision-making because they are capable of performing regression and classification tasks on complex data sets and solving non-trivial tasks. A frequently used example is the correct prediction of a XOr-logic gate output, which cannot be achieved by a linear decision boundary (Rosenblatt, 1958) but rather by an ANN with a hidden layer (Sprinkhuizen-Kuyper and Boers, 1996). Classification resembles the decision between discrete variables, e.g., “yes” or “no”, whereas a regression fits the provided data within the range of a continuous variable (Bourdès et al., 2010).

Attempts have been made to implement ANNs in NGS data analyses, as NGS is a well-established method in medicine, environmental microbiology, biotechnology and related fields. NGS generates a high number of sequencing reads of DNA and reverse-transcribed RNA. Therefore, an appropriate data format to supply the ANN with the information is essential to link the methods. One option is to use sequencing-derived or processed data, not the raw sequences themselves. An example is the correlation of the DNA methylation state with the age of a patient for forensic purposes (Vidaki et al., 2017). However, Nguyen et al. (2016) applied a convolutional neural network (CNN) to treat DNA sequences as a string input and store the position of the nucleotides in the sequence. In the CNN described by Umarov and Solovyev (2017) the promoter regions of eu- and prokaryotes were successfully predicted, whereas in Khawaldeh et al. (2017) DNA sequences were classified based on their taxonomy. In this study, 16S rRNA and rRNA gene amplicon sequencing was applied to generate data on microbial community assemblages. The sequence reads were phylogenetically clustered and taxonomically annotated by the SILVAngs pipeline Yilmaz et al. (2014). The resulting relative abundances of taxonomic clusters per sample were used as the input for the ANN (SI Fig. 1). The aim for the ANN was to automatically differentiate glyphosate-treated from untreated control communities. Furthermore, it was assumed that taxonomic clusters indicative of glyphosate pollution might be among those needed for the classification of the ANN, and their identification consequently could support monitoring efforts. Finally, both the robustness of the ANN setup and the amount of taxonomic information required for a reliable classification were investigated.

To our knowledge, NGS data on microbial community assemblages have not been used in an ANN-based classification. The fields of application of our findings are numerous and include the monitoring of specific events by classifying, e.g., contamination events or algal blooms as well as more generally fitting microbial community compositions via regression to, e.g., a salinity, temperature or temporal gradient.

## Material and Methods

### Laboratory & sampling

#### Laboratory experimental setup

Two 12-L (20×30×20 cm) microcosms comprising float glass and silicone glue were obtained from Rebie Aquaristik (Bielefeld, Germany). The microcosms were cleaned with EtOH (70%) and rinsed with sterile, filtered MilliQ water before they were filled with the sterilized substrates. Surface brackish water for inoculation was collected 2.5 km north of Warnemünde, Germany (54.199412, 12.042317). Five hundred millilitres of water was sterile filtered per GVWP filter (0.22 µm, Merck Millipore, Darmstadt, Germany) until the collected volume was filtered. The filters were immediately shock frozen in liquid nitrogen and stored at −80°C. Modified artificial brackish water (ABW) Bruns and Cypionka (2002) containing double the amount of KH_2_PO_4_ served as the substrate. A stock solution of 20 g casein hydrolysate (Merck, Darmstadt, Germany)⋅L^−1^ dissolved in MilliQ (Merck Millipore) served as the carbon and nitrogen source. The solution was sterile filtered (0.22 µm, Sartorius, Göttingen, Germany) and stored at 15°C. The casein hydrolysate was added to the ABW after autoclaving to a final concentration of 2.5 mL⋅L^−1^. Fire-dried quartz sand (0.1–0.4 mm, Quarzwerke, Frechen, Germany) was combusted for at least 4 h at 500°C in aluminium trays (Alcan, Brazil) and served as an artificial, carbon-free hard substrate. The microcosms were filled with 2 kg of quartz sand (~1.6 L) and 8 L of ABW. Combusted GF/F microfibre filters (Ø 47 mm, Whatman, Little Chalfont, UK) were placed into the hard substrate to provide easily collectible biofilm-overgrown material. Air pumps (2×200 L⋅h^−1^, 4 Watt, EHEIM GmbH, Deizisau, Germany) delivered sterile-filtered air (0.2 µm, Midisart 2000, Sartorius Stedim, Göttingen, Germany). Three thawed GVWP inoculum filters were cut in half, with one half placed in each microcosm overnight. ABW was refilled on days −56 and −34; the batch mode lasted from day −69 to day −31 to ensure that the bacteria formed biofilms on all surfaces, thereby preventing glyphosate adsorption (SI Fig. 1). Beginning on day −31, stable nutrient conditions were provided by changing the cultivation mode to a chemostat-like continuous culture to prevent substrate depletion and product accumulation. A peristaltic pump (Ismatec IPC 8, Cole Palmer, Wertheim, Germany) transported ABW from a sealed, autoclaved 5-L Schott bottle through clean, sterile tubing (Ø 1.02 mm (ID), silicone peroxide, Ismatec) at a flow rate of 0.37–0.38 mL⋅min^−1^ (537–548 mL⋅day^−1^) into the microcosms representing the water column. A second peristaltic pump with a flow rate of 0.33–0.34 mL⋅min^−1^ (475–489 mL⋅day^−1^) removed water from the opposite end of the microcosms such that excess volume was available for sampling. The 5-L Schott bottle was regularly exchanged together with the inlet tubing. On day 0, a pulse of sterile-filtered glyphosate (10 mg⋅L^−1^ final concentration) was added to the water column of the treatment microcosm and mixed by stirring.

#### Sampling procedure

Samples were taken for the determination of total cell counts and the glyphosate and aminomethylphosphonic acid (AMPA) concentration. One-hundred-millilitre water column samples were sterile filtered in three replicates; these filters were used for nucleic acid extraction. Samples for the DNA/RNA extraction were shock frozen in liquid nitrogen and stored at −80°C. Five-millilitre samples for the determination of glyphosate and AMPA concentrations were stored at 20°C without further treatment.

#### Nucleic acid extraction and sequencing

Nucleic acid extraction and DNA digestion were performed according to Bennke et al. (2018) for the filtered water samples. Biofilm samples were extracted using the phenol-chloroform method described in Weinbauer et al. (2002). cDNA synthesis was performed using 20 ng DNA-free total RNA as the input for the MultiScribe (Fisher Scientific GmbH, Germany) Reverse Transcriptase system with reverse primer 1492r (5’ TACGGYTACCTTGTTACGACTT (Lane, 1991)). Illumina amplicon sequencing was prepared as described in Bennke et al. (2018). The V3-V4 region on the 16S rRNA gene was targeted with the primer set 341f-805r (forward: CCTACGGGNGGCWGCAG, reverse: GACTACHVGGGTATCTAATCC (Herlemann et al., 2011)). Indexed amplicon libraries were pooled to a concentration of 4 µM. The PhiX control was spiked into the library pools at a concentration of 10%. Each final library pool (4 pM) was subjected to one of two consecutive individual paired-end sequencing runs for water column samples using 600 cycle V3 chemistry kits on an Illumina MiSeq. During the DNA amplicons run, 706 K⋅mm^−2^ clusters were sequenced; generating 17.6 million reads that passed the filter specifications. Over 70% of the sequencing and index reads were found to have a Qscore ≥30. During the cDNA amplicons run, 555 K⋅mm^−2^ clusters were sequenced. This generated 13.9 million reads passing filter specifications. Over 74% of the sequencing and index reads had a Qscore ≥30.

#### Bioinformatic and statistical analysis of amplicon data

Sequence data preparation for the SILVAngs pipeline was performed as described previously (Bennke et al., 2018). The SILVAngs pipeline dereplicated 100% identical sequences. Of the remaining unique 16S rRNA sequences, OTUs with a similarity threshold of 98% were selected. The representative sequence per OTU was taxonomically annotated using the ARB-SILVA database. Identical annotations for different OTUs were merged into clusters on the genus level; thus, the term “clusters” is used instead of OTUs. From the resulting taxonomy file containing reads per sample per cluster the clusters annotated as “No relative” were discarded. The relative abundance per cluster in % was calculated from the read fraction of the cluster of the library size of the sample. The unfiltered data set underwent no further quality check; for the filtered data set those with fewer than five reads were excluded.

The sequences were deposited in the NCBI database under BioProject ID PRJNA434253 and SRA accession SRP151042.

### Machine learning

#### Neural network architecture

The Java library for the ANN was Deep Learning for Java (DL4J), with N-dimensional arrays (Patterson and Gibson, 2017) and default values for most parameters. ANNs receive data via an input layer with a number of input nodes representing the dimensions of the input data; therefore, the input layer contained one neuron per feature (taxonomic cluster) in the respective data set, ranging from 1 to 687.

The signal is transported from one node to the next (layer), and the strength of the signal is altered by the weight of the connection, allowing for separation between more and less impactful nodes. The weights were Xavier initialized (default). Scaling of the signal between 0 and 1 was achieved using Softmax for output normalization. On each node or neuron, a threshold must be met by the incoming signal to activate the forwarding to the next layer. The neuron activation function was tanh (the default).

By employing further connected layers, the hidden layers, more complex interactions are enabled due to more combinations of input signals. Hidden layer 1 comprised 25 neurons, hidden layer 2 comprised 5 neurons and the output layer comprised 2 neurons, one for each class “treatment” and “control”. The output layer showed the aggregated result of the signals channelled through the preceding nodes. Each layer was fully connected to the next. The ANN had to be trained beforehand to classify the microbial community compositions. The option used was providing training data of a given format and amount as well as the expected classification, consequently the so-called supervised learning. Using backpropagation as an iterative learning process, the ANN adjusts the weights of the connections between the nodes to yield the provided classification. For this, the loss function was the negative log likelihood (Glorot and Bengio, 2010). Every experiment used 2000 learning cycles, with a learning rate of 0.1 for each repetition. Prior unknown data of the same format might then be classified by the trained ANN. To do so, the initial data set must be split into a training quota and a test quota. A third quota is required if, e.g., several ANN setups are to be compared and validated before the actual testing. It is therefore only feasible if sufficient data is available (Wu et al., 2013). As sequencing is still comparatively expensive, the amount of samples processed for this experiment was large but limited. Therefore, the data were split into training and test data sets, with the largest portion being training data.

#### Format of the main data sets

The taxonomy tables contained the relative abundance of a taxonomic cluster as the input feature for a given sample (SI Fig. 1). Each sample represented a unique combination of time point, nucleic acid, microcosm, habitat and - in case of the filtered data sets – technical replicate (2 or 3). All tables were designated either “treated” or “control”. The former denoted only samples in the glyphosate microcosms after glyphosate addition. Samples from the later-treated microcosm before the addition of glyphosate were labelled “control” as well (SI Fig. 1). Hence, data sets included more “control” than “treated” tables. Consequently, guessing “control” as classification would be correct for ~ 59% of the tables. We did not duplicate existing “treated” tables to generate a 1:1 ratio of “treated” and “control” as the deviation was not considered to be problematic.

Main data set 1 was the “unfiltered data” set, which consisted of all 687 clusters and the averaged technical replicates, resulting in 64 observations (38 x “control”, 26 x “treated”). Note that from the original experimental data containing information on water column and biofilm, the relative abundances of clusters within the biofilm samples were removed in this study at a later step but remained a feature of the taxonomy file format and were considered with an input node. Effectively, it represents setting all biofilm-originated clusters to 0. Therefore, the unfiltered data set contained 687 taxonomic clusters with 213 being biofilm-originated, resulting in 474 clusters different from 0. One observation was randomly selected to test the classification performance of the ANN; the remaining tables comprised the training data.

Main data set 2 was labelled “filtered data”. The tables were filtered before the relative abundances were calculated by removing clusters with less than five counts in a sample. Additionally, the replications were not averaged but rather used as separate observations, yielding 187 observations (111 x “control”, 76 x “treated”). Consequently, the filtered data set contained 360 taxonomic clusters with 86 clusters being biofilm-originated, resulting in 274 clusters different from 0. One observation was randomly selected for testing; the remaining tables comprised the training data. Excluded were the remaining tables of the replicate as they were very similar to the test observation. Which observations (tables or samples) and which features (taxonomic clusters) were additionally used in the various classification setups is illustrated in SI Fig. 2.

#### Test ANN classification

The general applicability of ANNs in classifying community composition data was tested using the unfiltered data set as the input. The classification was repeated 2000 times.

#### Identifying clusters present in a successful classification

To identify the taxonomic clusters participating in successful classifications, a subsetting approach was applied. For the unfiltered data set, 30 clusters were chosen randomly, and the network was trained with those 30 clusters. For each subset, the classification was repeated 256 times (so with 64 tables each covering 4 times the test table), and the number of correct classifications and the chosen clusters were documented. The order of averages stabilized after ~1000 subsets, and the process was stopped at 1220 subsets. Each subset classification required approximately 1 hour on a virtual machine with 4 CPU cores and 16 GB RAM. Comparably for the filtered data set, the top 10 ranking clusters were calculated by the random subsetting approach based on a sample size of 20 clusters. Subsetting was performed 1066 times, and on each subset 1000 classification runs were performed. Four specific clusters appearing in the unfiltered and filtered subsets were compared regarding their participation in correct classifications.

#### Determine required feature amount for classification

The previous experiment ranked each cluster based on the number of times it was present in successful classification subsets. From this order, the 10 top-ranked clusters were selected to determine whether a classification was possible with a significantly reduced number of features. Furthermore, the limitations of cluster reduction were explored by sequentially removing the lowest-ranked cluster of the top 10.

#### Determine required number of observations for classification

To examine whether RNA or DNA data alone were sufficient input for classification and which is better suited, the unfiltered data set was split into 32 DNA and 32 RNA tables, with 31 tables used for training and the remainder for testing. We conducted 13568 repetitions for classification based on DNA and 2048 repetitions based on RNA. Additionally, to investigate the required sampling resolution, half of the time points from the unfiltered data set were evenly distributed and removed from the data set, and the classification was tested 6656 times.

#### Other machine learning algorithms tested

In the process of analysing the community assemblage data, other machine learning algorithms were tested provided by the Weka toolkit (Hall et al., 2009). Machine learning approaches included Decision Table and Random Forests algorithms and their ANNs. None of the approaches provided correct classifications with standard parameters, and the computation times of the respective ANNs were excessive.

## Results

### ANN identifies glyphosate-treated microbial communities

The ANN achieved 1905 correct classifications out of 2000 repetitions (95.25%) using the unfiltered data. A classification based purely on guessing would have been correct ~ 59% of the time due to the uneven distribution of the “treated” and “control” observations. If only the microbial community assemblages from the two microcosms were distinguished rather than those which actually experienced contact with glyphosate, at best a 90.625% correct classification could theoretically have been reached. It is therefore generally possible to use an ANN to separate community composition data.

### Identification of clusters present in successful classifications

To understand which clusters participate in correct classification and hence are possibly important, subsets comprising randomly chosen clusters were tested. For the unfiltered data set, the subset size was 30 clusters, and 1220 subsets were generated. Of 256 classification repetitions for each subset, the number of correct classification ranged from 30.1 - 98.4% (Fig. 1). For the filtered data set, 1066 subsets with a size of 20 clusters were selected, and on each subset 1000 classifications were performed, with the correct classification per subset ranging from 36.7 – 99.7%. The range is comparable; the distribution of the classification, however, displayed an increase in valuable subsets for the filtered data set. Fifty percent of the subsets of unfiltered data achieved a classification rate centred around the guessing level, and only a small fraction of subsets was above the 90.625% threshold compared with a significantly reduced fraction for the filtered data set at guessing level and increased fraction at all higher classification rates, especially around the critical 90.625% threshold. Furthermore, an increased average of the classifications was observed as well as reduced computational efforts, as the distribution of classifications stabilized after 300 subsets compared with 1000 subsets for the unfiltered data set.

**Figure 1.**
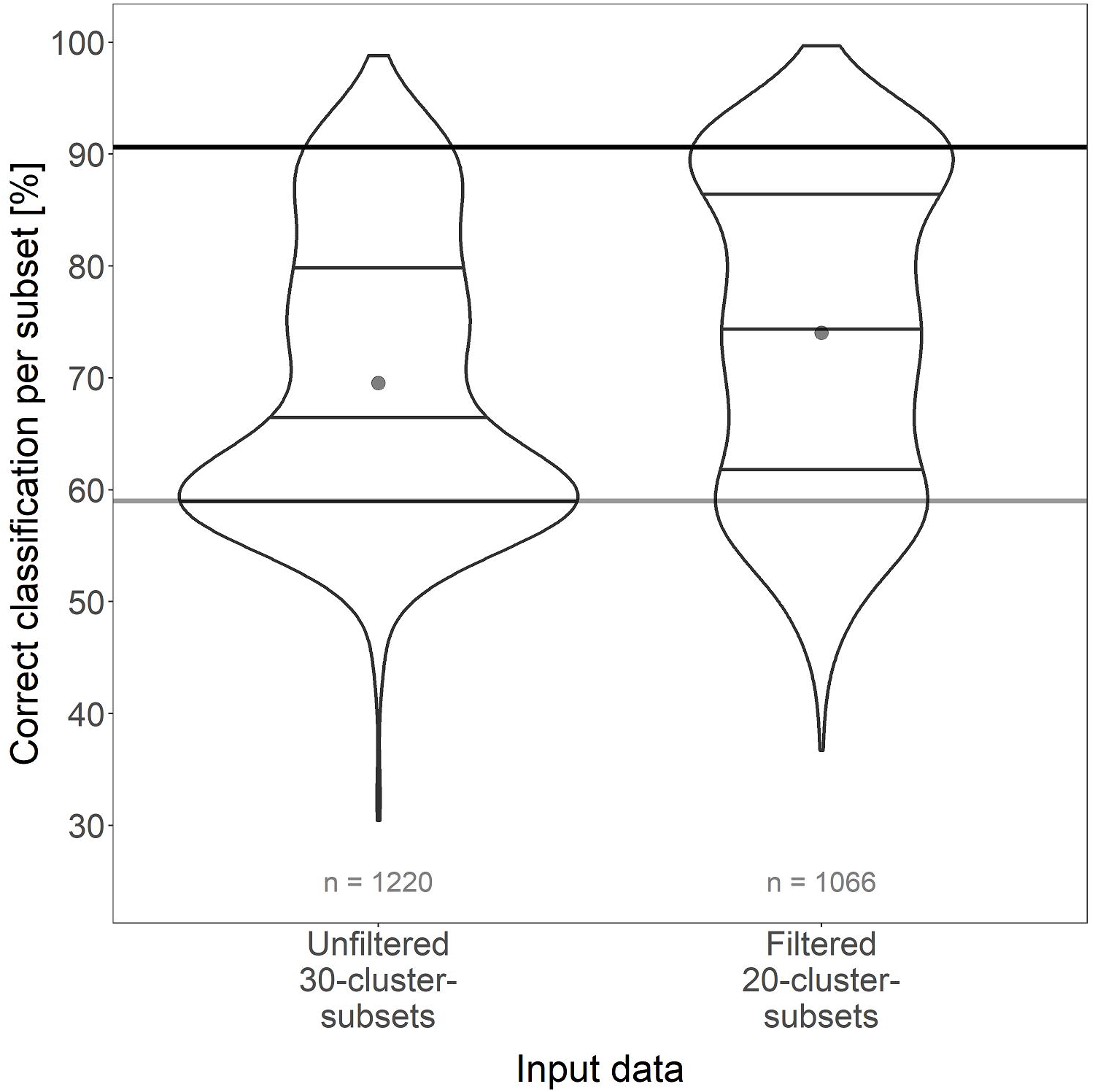
Violin plots of correct classification rates by random subsets of size 30 for the unfiltered data set and size 20 for the filtered data set, respectively. *n* is the number of subsets that were generated for the respective data set. The dot represents the average, the three horizontal lines within the violin plot depicture the 25%, 50% and 75% quantiles. The horizontal bar at 59% displays the classification achievable by pure guessing, the upper bar marks the threshold for a classification which both separates the microcosms and before and after glyphosate addition. A shift from many subsets classifying around the guessing level for the unfiltered data set to improved classification rates in the filtered data set is shown.

The ranking, based on the average classification from all subsets a cluster was part of, revealed which clusters were frequently part of correctly classified subsets. The classification of subsets containing the clusters *Massilia* spp., *Parvibaculum* spp., *Gallaecimonas* spp. and *Limnohabitans* spp. were compared (Fig. 2; relative abundances in SI Fig. 3 a, b, l and r). They appeared in both data sets. In particular, *Gallaecimonas* spp. and *Parvibaculum* spp. displayed a distinct increased relative abundance following the glyphosate addition, which could be useful for the classification by the ANN. *Massilia* spp. and *Limnohabitans* spp. are clusters for which the temporal abundance course did not reveal a response to glyphosate but rather differed between the two microcosms. *Massilia* spp. in both data sets were part of the well-performing subsets and identified as a very valuable cluster for classification in the setup described below.

**Figure 2.**
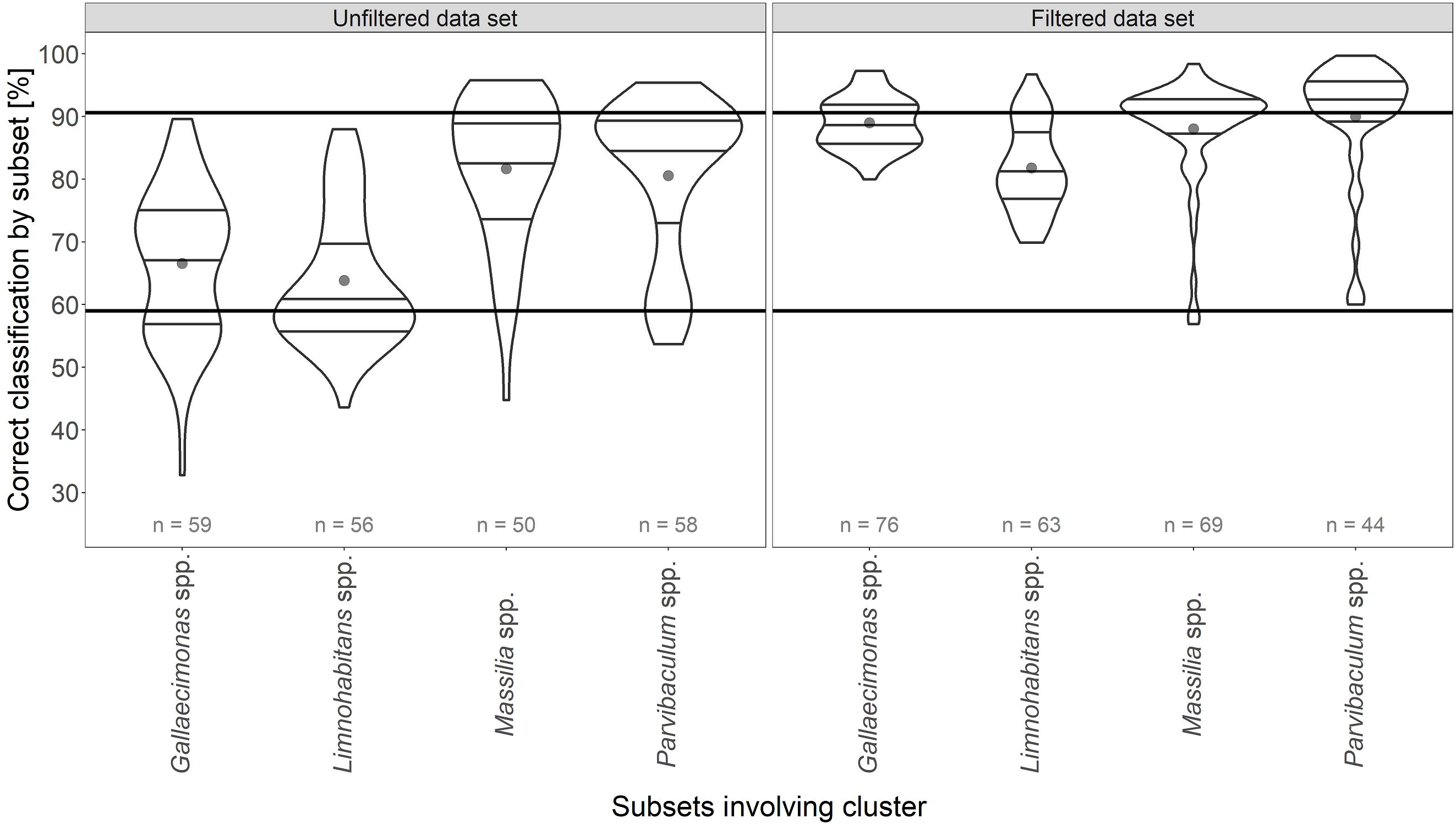
Violin plots of correct classification by subsets containing specific taxonomic clusters for both data sets. *n* is the number of subsets that included the respective cluster. The dot represents the average classification, the three horizontal lines within the violin plot depicture the 25%, 50% and 75% quantiles. The horizontal bar at 59% displays the classification achievable by pure guessing, the upper bar marks the threshold for a classification which both separates the microcosms and before and after glyphosate addition. The filtering step improved the ANN’s performance by reducing the range and frequency of less good classifications towards a higher number of better classifications. *Gallaecimonas* spp. containing subsets drastically improved classification rates.

As shown in Fig. 2, subsets containing *Gallaecimonas* spp. and *Limnohabitans* spp. did not perform well on the unfiltered data set, and the classification rates for *Gallaecimonas* spp. containing subsets ranged from 34 - 89%. This changed in particular for subsets of the filtered data set containing *Gallaecimonas* spp., as a high fraction of the subsets classified approximately 90% correctly; however, subsets with *Limnohabitans* spp. improved, too. *Parvibaculum* spp. containing subsets showed a good performance in both data sets but improved still in the filtered data as they reached the upper threshold for 50% of the subsets, similar to subsets containing *Massilia* spp. For both data sets it should be noted that performing the actual classification with an ANN trained on random subsets is not effective. The averaged classification rate over all generated subsets from filtered data is 74.1% and 69.5% from unfiltered data.

### Exploring the limits of the required cluster features and assembling a highly indicative selection

The random subsets were expected to be unable to perform a sufficient classification contrary to the full data set. The results indicate that some clusters are more decisive than others. Thus, the 10 clusters with the best average classification rate from the subsetting approaches were selected as the sole training data for the ANN. This yielded a classification rate of 94.4% for the unfiltered data and 95.8% for the filtered data (Fig. 3). The two top 10 cluster lists contained different clusters for the respective data set (Tab. 1) which might be caused by the removal of certain clusters due to their low abundance during the filtering step. The maximal relative abundance per top 10 clusters ranged from 0.07 % (*Dokdonella* spp., only unfiltered data) and over 0.76 % (*Parvibaculum* spp., both sets) to 9.27 % (*Gallaecimonas* spp., both sets). Essentially, both top 10 data sets yielded a classification as good as the full data set (95.25%). Further reducing the number of features revealed that using at least the six best-ranked clusters as the input was required to yield a classification rate > 90.625% for filtered data, whereas the unfiltered data sets kept meeting the threshold using as few as two clusters. A further stepwise reduction of the filtered data to only the top two clusters lowered the classification rate to 88.8%. Interestingly, the classification rate with unfiltered data decreased to near the guessing level (63.5%) when only the best ranked cluster was used (*Massilia* spp., Tab. 1; SI Fig. 3 a). *Massilia* spp. comprised a cluster only abundant in the control microcosm. The second best unfiltered cluster was *Parvibaculum* spp., and it was determined to be the best-ranked cluster for the filtered data set and, in contrast, performed well on its own (91.7%). The relative abundance of *Parvibaculum* spp. was different between both microcosms as well as before and after the glyphosate addition (SI Fig. 3 a). It was also observed that 2 clusters yielded a better classification than 3 or 4 for filtered data, and the decrease was not linear for the unfiltered data. *Parvibaculum* spp. were tested as a sole feature for both data sets and yielded a significant increase in the filtered data set.

**Table 1.** Listing and comparison of all taxonomic clusters revealed by random subsetting or bioinformatic analysis applying R package DESeq2. Includes the rank, maximum abundance per DNA or cDNA targeted approach and the *p* value, if available.

| Genus | Family | Order | Class | Phylum | Unfiltered<br>data top 10<br>rank | Filtered<br>data top<br>10 rank | DESeq2 Wald<br>test adjusted<br>*p*-value | Plot ID in<br>SI Figure<br>3 | maximal relative<br>abundance for<br>DNA or RNA [%] |
| --- | --- | --- | --- | --- | --- | --- | --- | --- | --- |
| *Massilia* | *Oxalobacteraceae* | *Burkholderiales* | *Betaproteobacteria* | *Proteobacteria* | 1 | 4 | *** | a) | 7.6 |
| *Parvibaculum* | *Rhodobiaceae* | *Rhizobiales* | *Alphaproteobacteria* | *Proteobacteria* | 2 | 1 | *** | b) | 0.76 |
| *Dokdonella* | *Xanthomonadaceae* | *Xanthomonadales* | *Gammaproteobacteria* | *Proteobacteria* | 3 |  |  | c) | 0.07 |
| *Reyranella* | *Rhodospirillales* | *Rhodospirillales* | *Alphaproteobacteria* | *Proteobacteria* | 4 |  |  | d) | 0.09 |
|  |  | B38 | *Gammaproteobacteria* | *Proteobacteria* | 5 |  | *** | e) | 0.12 |
| *Loktanella* | *Rhodobacteraceae* | *Rhodobacterales* | *Alphaproteobacteria* | *Proteobacteria* | 6 |  |  | f) | 0.48 |
| *Caulobacter* | *Caulobacteraceae* | *Caulobacterales* | *Alphaproteobacteria* | *Proteobacteria* | 7 | 7 |  | g) | 0.37 |
| *Aminobacter* | *Phyllobacteriaceae* | *Rhizobiales* | *Alphaproteobacteria* | *Proteobacteria* | 8 |  |  | h) | 0.15 |
| *Nesiotobacter* | *Rhodobacteraceae* | *Rhodobacterales* | *Alphaproteobacteria* | *Proteobacteria* | 9 |  |  | i) | 0 |
| *Idiomarina* | *Idiomarinaceae* | *Alteromonadales* | *Gammaproteobacteria* | *Proteobacteria* | 10 |  | * | j) | 0.62 |
| *Hyphomonas* | *Hyphomonadaceae* | *Caulobacterales* | *Alphaproteobacteria* | *Proteobacteria* |  | 2 | *** | k) | 0.66 |
| *Gallaecimonas* | Unknown Family | *Gammaproteobacteria* | *Gammaproteobacteria* | *Proteobacteria* |  | 3 | *** | l) | 9.27 |
| *Thalassobaculum* | *Rhodospirillaceae* | *Rhodospirillales* | *Alphaproteobacteria* | *Proteobacteria* |  | 5 | *** | m) | 7.66 |
| *Sphingopyxis* | *Sphingomonadaceae* | *Sphingomonadales* | *Alphaproteobacteria* | *Proteobacteria* |  | 6 |  | n) | 2.75 |
| *Rhizobium* | *Rhizobiaceae* | *Rhizobiales* | *Alphaproteobacteria* | *Proteobacteria* |  | 8 | *** | o) | 6.22 |
| *Brevundimonas* | *Caulobacteraceae* | *Caulobacterales* | *Alphaproteobacteria* | *Proteobacteria* |  | 9 |  | p) | 0.37 |
| *Sphingomonas* | *Sphingomonadaceae* | *Sphingomonadales* | *Alphaproteobacteria* | *Proteobacteria* |  | 10 |  | q) | 1.75 |
| *Limnohabitans* | *Comamonadaceae* | *Burkholderiales* | *Betaproteobacteria* | *Proteobacteria* |  |  |  | r) | 20.97 |

Column I: excerpt of statistically significant abundant taxonomic clusters after addition of glyphosate in the treatment microcosm. Statistical significance (\*\*\* < 0.001. \*\* < 0.01. \* < 0.1) is based on Benjamini-Hochberg corrected *p*-values . tested with Wald test using R package DESeq2

**Figure 3.**
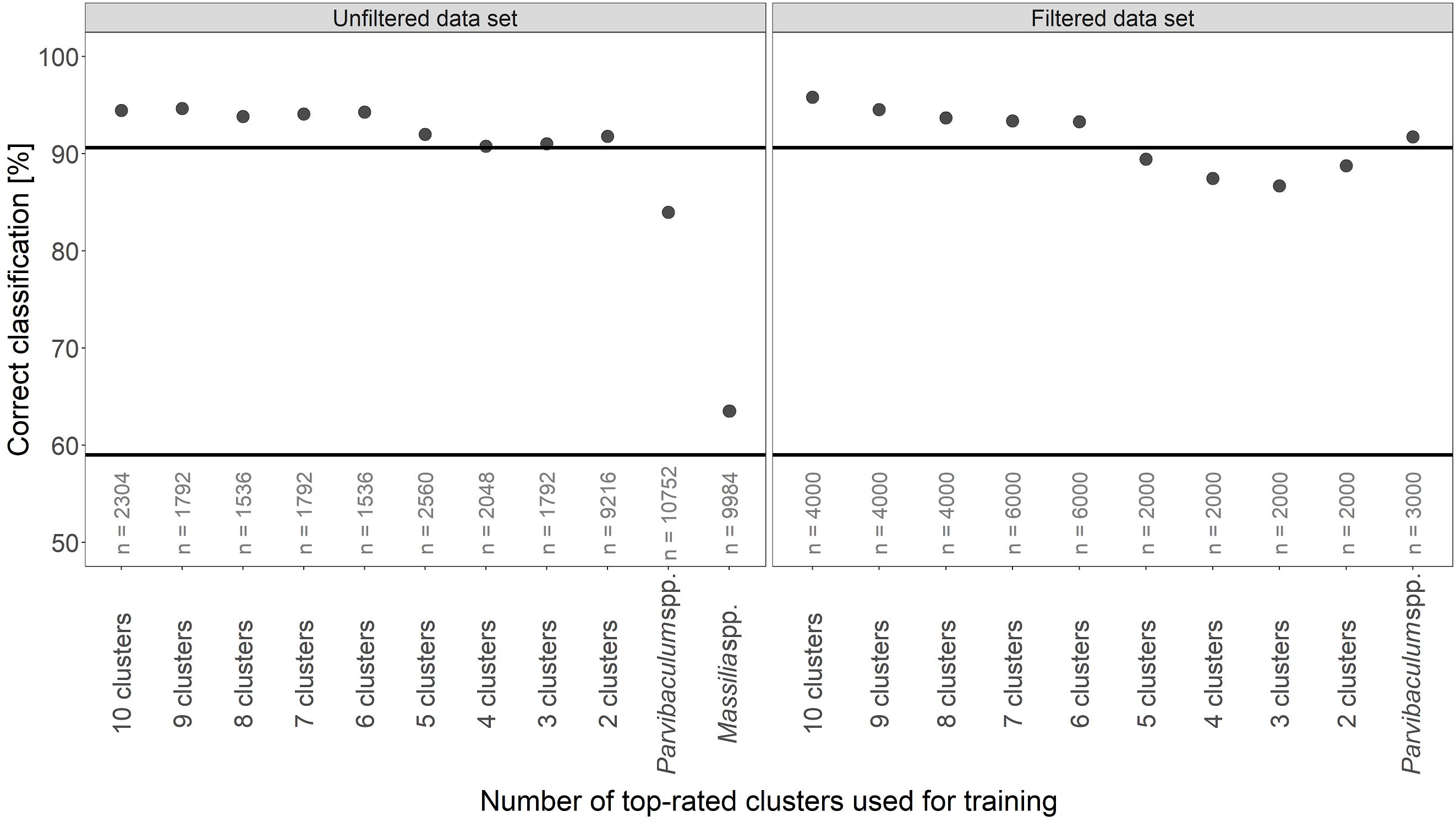
Classification rates achieved by using a top ranked selection of clusters. *n* is the number of classifications performed with the respective clusters. The horizontal bar at 59% displays the classification achievable by pure guessing, the upper bar marks the threshold for a classification which both separates the microcosms and before and after glyphosate addition. Information on 10 clusters is sufficient to classify as well as using the full data set. Removing one cluster at a time from the input did not result in a linear decrease. Depending on the cluster, one can provide sufficient data (*Parvibaculum* spp.). In that case, the classification was improved by the filtering step.

### Comparing the use of DNA-*vs.* RNA-derived data

After investigating the number of features used for classification, these approaches targeted the number of observations required. The unfiltered data set was used. The classification rate decreased to 82.2% if only DNA data was used and to 84.8% for the RNA-derived data (SI Fig. 4). Both values were within the range needed to distinguish between the two microcosms regardless of the glyphosate addition. Excluding half of the sampling time points resulted in a classification of 82.1%.

## Discussion

Information collected over 16 time points from a microbial community assemblage obtained in a lab microcosm experiment in which glyphosate was applied as a disturbant was used to train an ANN for the classification of treated and control communities. Glyphosate is not considered as strong a microbial stressor as, for example, toxic or antibiotic substances and, in fact, can be utilized as a nutrient or energy source by many microbes such that positively reacting clusters could be used for classification (Hove-Jensen et al., 2014; McGrath et al., 1997). The artificial neural network successfully distinguished between treated and untreated communities and demonstrated the general feasibility of the combined NGS-ANN approach. In particular, the ANN was required to separate community compositions of two independent microcosms with slightly different assemblages from the beginning of the experiment, which was easily achieved by standard ordination methods such as non-metric multidimensional scaling (SI Fig. 1). The experimental design also demanded the ANN to identify traits present in the control-labelled samples from both microcosms and to separate those from the treated samples, which it successfully accomplished.

### A statistical approach to identifying decision-important clusters improved with fewer features

The subsequently applied random subsetting method was developed because a systematic approach was not considered feasible for this study; any expedient selection of factors resulted in combinatorial explosion. The subsetting led to two important conclusions:

1. It was possible to stochastically identify and rank input features. This involved the identification of important taxonomic clusters to differentiate between the samples. Therefore, it could help determine indicator candidates for environmental monitoring purposes.
2. The required number of features could be significantly reduced, as demonstrated by the equally successful classification by only the top 10 ranked clusters of each data set. Using less than approximately 10 clusters might result in a loss of required information (Fig. 3). This was shown by the non-linear decrease in classification rates. This indicated that each cluster may contribute a certain dimension of information; the conducted removal of clusters was based on their average classification within a subset, which may not reflect the value of these pieces of information. To conclude, it is presumably not worth the reduced computational costs to base classification on a single-digit number of features.

The filtering of the data shortened the time needed for the subset ranking to stabilize, which might also depend on the size of the subset. Several comparisons between the two data sets, e.g., of subsets (Fig. 1) containing specific clusters (Fig. 2) and the classification trained solely on *Parvibaculum* spp. abundance (Fig. 3), proved a significant increase in the contribution to a successful classification if the data was filtered from low abundant clusters. We encourage the application of a filtering step on microbial community composition data sets for similar approaches. Solely random subsampling led on average to unsatisfactory classification results, which indicated that each participating cluster may also contribute its information in a misleading way, e.g., clusters that were unresponsive to glyphosate or empty clusters. However, a few subsets in both data sets breached the classification threshold of 90.625%. In Fig. 2, the contribution of certain clusters towards the classification performance of their subset is displayed. It can be assumed that evenly distributed classifications ranging between many percentages (*Gallaecimonas* spp. in unfiltered data) indicate that the cluster within this subset is not a dominant contributor of information; hence, the classification success rather depends on the other members of the subset. A distinct range of classification within a subset (filtered data, *Parvibaculum* spp.) might rather indicate a decisive cluster of a subset. The clusters *Parvibaculum* spp. and *Massilia* spp. were part of both top 10 lists. How *Massilia* spp. supported a classification, while being present in only one microcosm, is a matter of speculation. It may be that the abundance of *Massilia* spp. separates the microcosms while another cluster contributes the information needed to distinguish between before and after the glyphosate addition (SI Fig. 3 a, b). This is supported by the sharp fall in classification rate when *Massilia* spp. were used as a single input feature, whereas the ANN solely trained on *Parvibaculum* spp. succeeded. The bioclimatic model of Larsen et al. (2012) predicted bacterial community assemblages on order level. It incorporated 16S rRNA gene pyrosequencing data in a preceding data analysis step before applying an ANN to spatially and temporally extrapolate microbial diversity. Their findings suggested that the strength of the ANNs is to combine the information on abundances of multiple taxa. Another interesting finding was the absence of an abundance of *Nesiotobacter* spp., a top 10 cluster (SI Fig. 3 i). Its appearance in only the unfiltered data suggests that it represented the many “empty” clusters that were also part of the “treated” tables. It could also be coincidentally part of the well-performing subsets.

### More observations should be generated

In general, more complex data sets require more complex models to explain the data. If the data sets additionally are noisy, characterized by a larger variance, even more training data and - more specific - more observations to adjust the weights of the ANN are necessary. This was demonstrated in experiments with decreased numbers of taxonomic tables. The use of only RNA- or DNA-derived data as well as only half of the time points reduced the classification rate to below 90.625% (SI Fig. 4). It is also possible that the decrease was due to the small sample size. Therefore, an accurate determination of whether DNA- or RNA-derived data is better suited and which temporal resolution is required for the training data was not possible. However, these findings indicate that the present number of samples is close to the limit for maintaining a correct classification. Fortunately, if such an ANN could be implemented in monitoring programmes, additional data would be generated at each monitoring event, which can be progressively included into the model such that the observations-to-features ratio is continuously improved.

### The outcome of the ANN was confirmed by bioinformatic analysis

The samples from the same glyphosate incubation experiment were also examined with bioinformatics tools, guided by slightly different hypotheses. From the 17 unique clusters in the unfiltered and filtered top 10 clusters (Tab. 1), which were identified by random subsetting approaches, eight were also identified by the R package DESeq2, which tests for statistically significant differences in abundance and was developed for NGS data (Love et al., 2014). It was applied to compare the cluster abundances before and after the glyphosate pulse. Subjecting the DESeq2 input to a filtering step excluded some of the clusters identified by the ANN. This step was thought to improve the reliability of statistics. The data suggests that the ANN can profit from low-abundant clusters as well (Tab 1). A combination of traditional bioinformatic or molecular ecology approaches and ANN technologies seems practical.

### Further steps in the application of ANN with NGS

While the results presented by this study are promising, the community assemblage data were still low-dimensional, containing information on the relative abundance per cluster, time point, glyphosate treatment, technical replication, and nucleic acid analysed. The samples were treated as independent observations. To make use of the capacity and potential of machine learning technologies, various aspects can be targeted for improvement. For example, the number of dimensions could be increased by adding meta data, often available from standardized monitoring campaigns, to the input. Temporal or spatial information should be included. At this step, a shift from classification to regression could be appropriate.

However, the first aim here was a robust ANN that can achieve a correct classification based only on sequencing-derived data. OTUs, or in this case, clusters, inherit a vast amount of functions, and their predictive ability is therefore limited to specific scenarios and environments. This can be leveraged by using available 16S rRNA gene amplicon data sets, e.g., from the Baltic Sea with known meta data. ANNs could be trained on these data as “standard”, and if a new sample is not classified as “standard”, it should be investigated to identify the reason for the deviation. The next logical step would be to use data from metagenomic and metatranscriptomic sequencing. This would resemble a function and not phylogeny-targeted approach, in which case the features would be the abundance of genes or their transcripts. The general principle was already demonstrated by Lin et al. (2017) who, employing a CNN, improved the assignment of single-cell RNAseq reads to their cell types of origin.

Although our suggested approach would necessitate more training data, the approach is considered feasible as sequencing costs are decreasing, and many suited data sets for training, validation and testing are publicly available.

## Acknowledgements and funding

We greatly appreciate comments from John Edward Boon, Jr. on a previous version of the manuscript that significantly improved the quality. The work was funded by the German national BMBF project “Sektorale Verwertung” (01IO1448). It also resulted from the EU-BONUS BLUEPRINT project and was supported by BONUS (Art 185), funded jointly by the EU and the German Federal Ministry of Education and Research (BMBF; 03F0679A).

## Supporting information

**SI Figure 1.** A timeline of the laboratory work flow followed by wet lab downstream processing, MiSeq sequencing and bioinformatic analysis. The taxonomic annotation was performed by the SILVAngs pipeline and the NMDS ordination plot was generated using the metaMDS function from R package vegan based on Bray Curtis dissimilarity. Red labeled samples experienced contact with glyphosate. The ascending alpha gradient indicates passing time.

**SI Figure 2.** A flow chart displaying the various approaches to detect the limits of reasonable classification by the ANN by reducing the amount of features and observations. Steps on unfiltered data are marked in red, filtered in green.

**SI Figure 3.** All top 10 ranked clusters (Table 3) from the filtered and unfiltered data plus *Limnohabitans* spp. were displayed based on their relative abundance with DNA- and RNA-derived data for both microcosms. The technical replicates are shown as dots, the mean as line. DNA is shown as continuous and RNA as broken line. The black vertical line demarks the addition of glyphosate. Due to abundance differences in orders of magnitude, the y scale is adjusted for each plot:

a) *Massilia* spp.; b) *Parvibaculum* spp.; c) *Dokdonella* spp.; d) *Reyranella* spp.; e) B38/*Gammaproteobacteria*; f) *Loktanella* spp.; g) *Caulobacter* spp.; h) *Aminobacter* spp.; i) *Nesiotobacter* spp.; j) *Idiomarina* spp.; k) *Hyphomonas* spp.; l) *Gallaecimonas* spp.; m) *Thalassobaculum* spp.; n) *Sphingopyxis* spp.; o) *Rhizobium* spp.; p) *Brevundimonas* spp.; q) *Sphingomonas* spp.; r) *Limnohabitans* spp.

**SI Figure 4.** Classification rates after removal of observations. *n* is the number of classifications performed with the respective setup. The horizontal bar at 59% displays the classification achievable by pure guessing, the upper bar marks the threshold for a classification which both separates the microcosms and before and after glyphosate addition. None of the setups was able to reach the upper threshold.

