## Supplementary figures and images for "An artificial neural network identifies glyphosate-impacted brackish communities based on 16S rRNA amplicon MiSeq read counts"

### Supplemental Figure 1

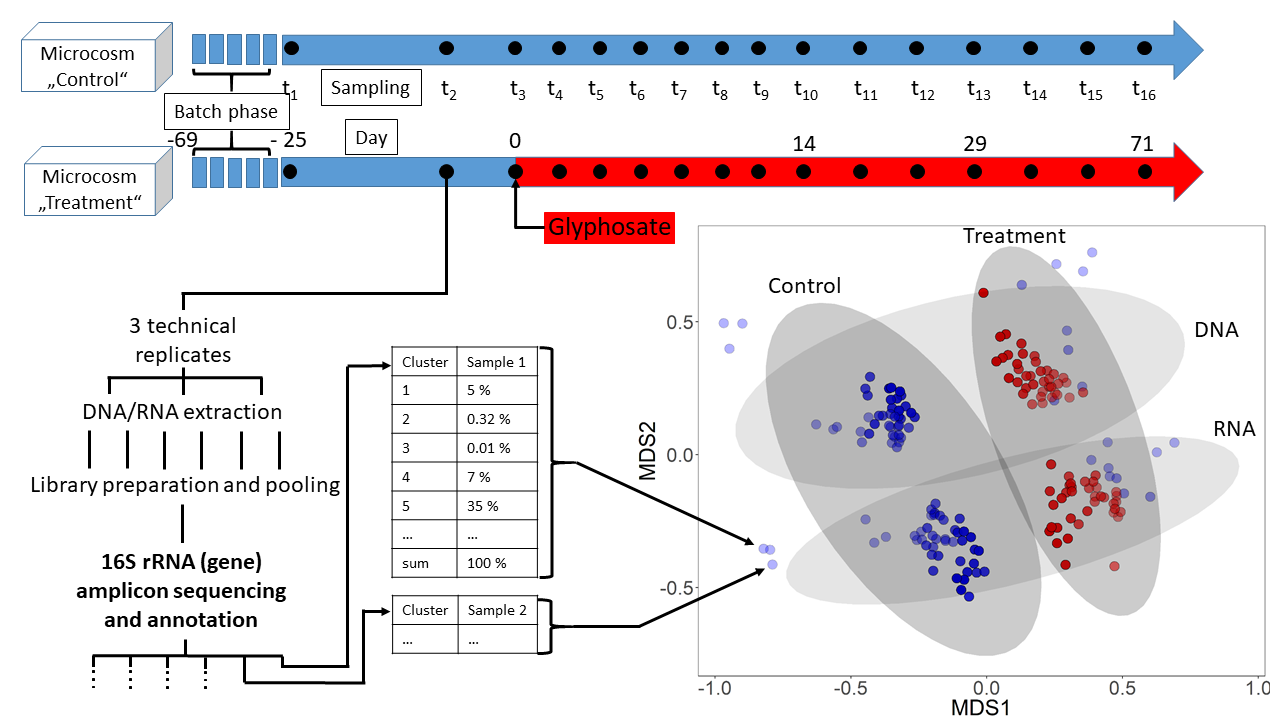

### Supplemental Figure 2

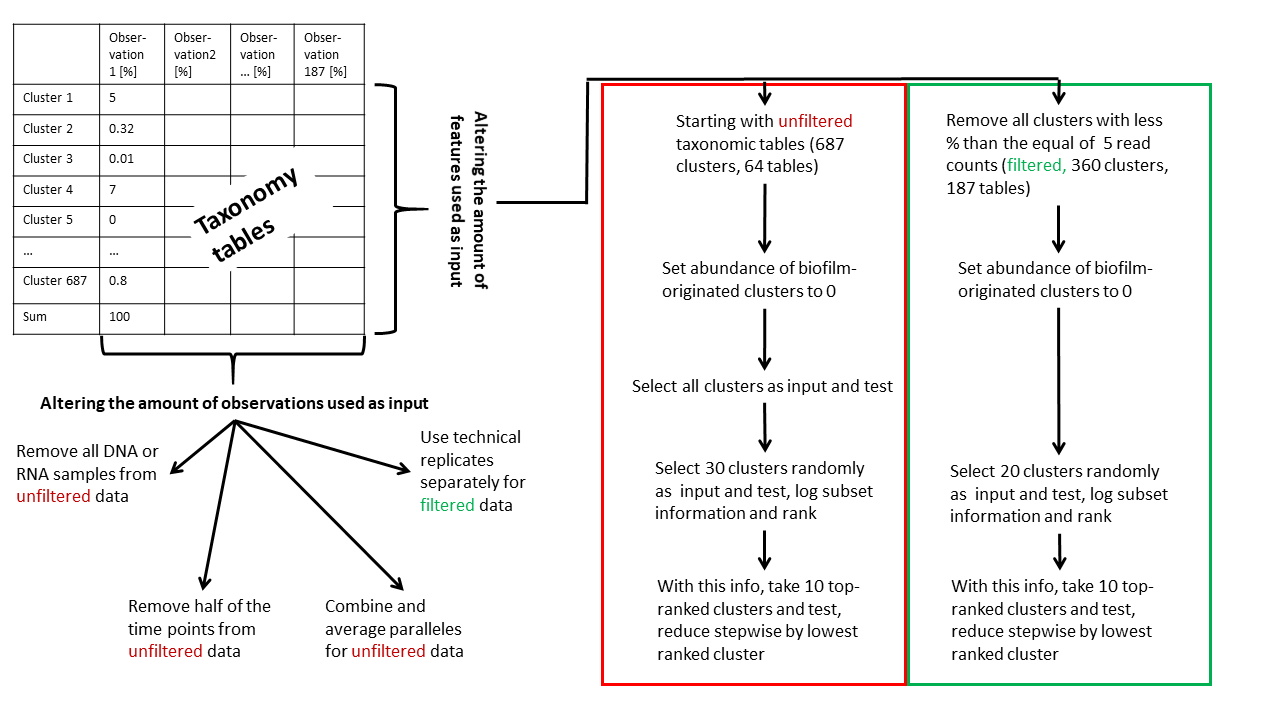

### Supplemental Figure 3

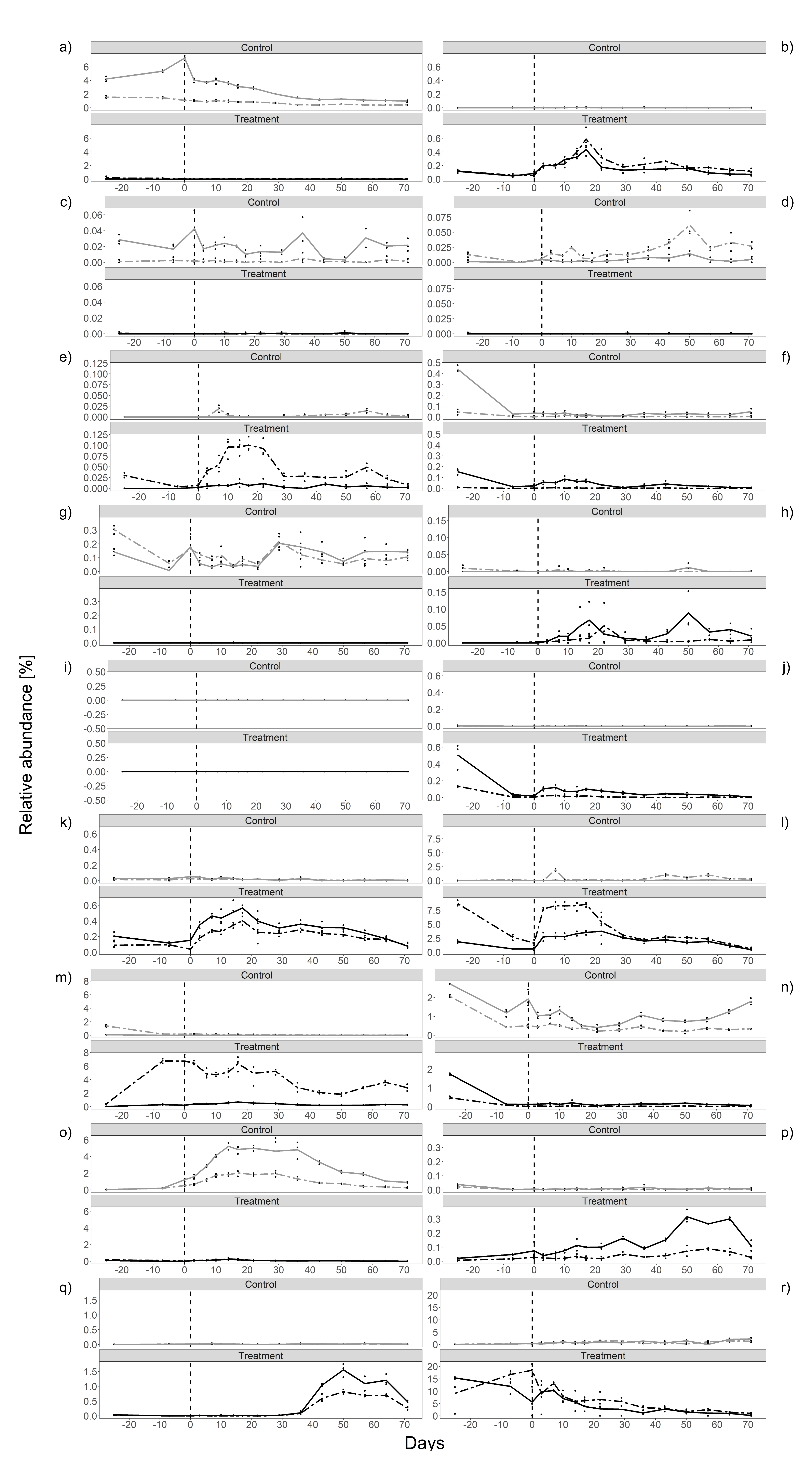

### Supplemental Figure 4

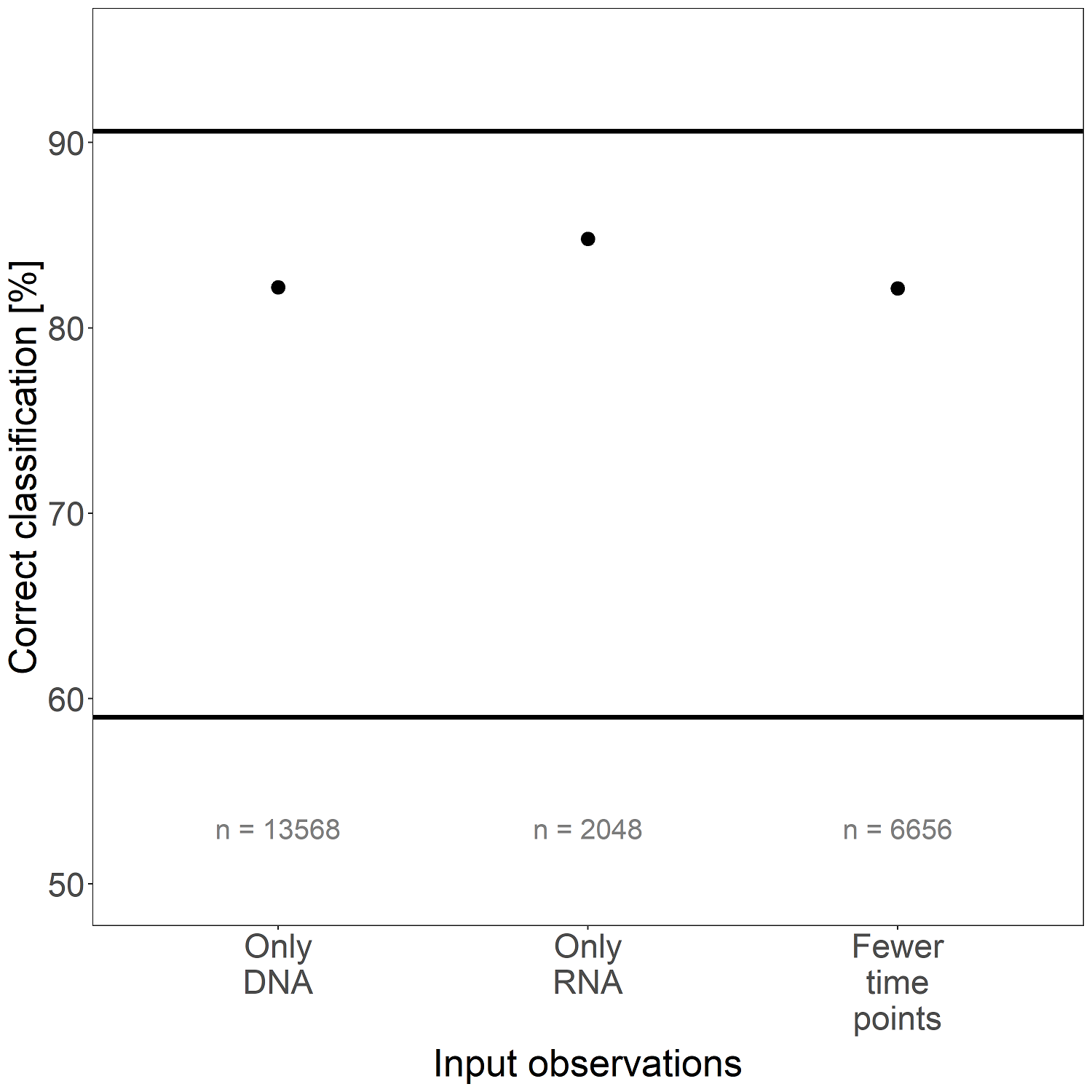
